# *Mycobacterium tuberculosis* m^4^C DNA methyltransferase Rv3204 promotes mycobacteria survival under oxidative stress

**DOI:** 10.1101/2025.08.26.672393

**Authors:** Abulimiti Abudukadier, Zhang Qiao, Lei Zhang, Chen Haiqi, Jingjing Niu, Li Peibo, Gong Zhen, Jianping Xie

**Affiliations:** Institute of Modern Biopharmaceuticals, School of Life Sciences, Southwest University, Chongqing, 400715 China; Department of Clinical Laboratory, theSecond Affiliated Hospital of Anhui Medical University, Hefei, Anhui, 230601, China; Chongqing Public Health Medical Center, 400030 China

**Keywords:** *Mycobacterium tuberculosis*, DNA methyltransferase, ROS, m4C

## Abstract

Reactive oxygen species (ROS) inflict cellular damage yet are pivotal mediators of signaling pathways. *Mycobacterium tuberculosis* (Mtb), the causative agent of tuberculosis (TB), persists as a major global health threat, partly due to its capacity to neutralize host-derived ROS toxicity. While DNA methylation deficiency is known to attenuate Mtb virulence, the specific role of DNA methyltransferases in mycobacterial survival under oxidative stress remains poorly defined. Here, we demonstrate that the mycobacterial protein Rv3204 functions as an N^4^-methylcytosine (m^4^C) DNA methyltransferase. Deletion of the *Rv3204* homolog (*Ms_1939*) in *Mycobacterium smegmatis* significantly impaired bacterial survival upon rifampicin exposure. This phenotype was associated with heightened intracellular ROS accumulation and a failure to upregulate transcription of ROS detoxification genes. Furthermore, the mutant exhibited downregulated expression of DNA repair genes and increased susceptibility to fluoroquinolone antibiotics (norfloxacin, ofloxacin). Crucially, the *Ms_1939* deletion strain displayed elevated levels of DNA damage. To our knowledge, this is the first study establishing a direct link between an m^4^C DNA methyltransferase and ROS homeostasis in mycobacteria. Our findings identify Rv3204 as a potential novel therapeutic target for modulating ROS sensitivity in *M. tuberculosis*.

**Importance:** *Mycobacterium tuberculosis*, the causative agent of tuberculosis, remains a major global health threat due in part to its ability to withstand host-derived oxidative stress. This study identifies Rv3204 as a novel N4-methylcytosine DNA methyltransferase essential for mycobacterial survival under oxidative stress. We demonstrate that deletion of *Rv3204* homolog in *M. smegmatis* leads to elevated intracellular ROS, impaired transcriptional activation of antioxidant and DNA repair genes, increased DNA damage, and heightened susceptibility to fluoroquinolone antibiotics. Our findings establish a critical link between epigenetic regulation via m^4^C methylation and ROS homeostasis in mycobacteria—a relationship previously unexplored. This work not only advances our understanding of bacterial epigenetic mechanisms in stress adaptation but also positions Rv3204 as a promising target for novel anti-tuberculosis strategies aimed at disrupting redox balance and enhancing antibiotic efficacy.

**DATA summary:** Raw data files have been deposited in the National Center for Biotechnology Information (NCBI) Gene Expression Omnibus (GEO) under accession number GSE139646 (https://www.ncbi.nlm.nih.gov/geo/query/acc.cgi?acc=GSE139646).

## Introduction

Reactive oxygen species (ROS) including hydrogen peroxide (H_2_O_2_), hydroxyl radicals (-OH), superoxide ions(O^2-^), are metabolic byproducts of aerobic respiration and responsible for maintaining cellular redox homeostasis. This balance, maintained by antioxidant enzymes, reduced thiols, and NADP(H) cofactors, is critical for both host antimicrobial defenses and pathogen survival. ROS can damage bacterial DNA and potentiate antibiotic efficacy through oxidative DNA lesions ^[1]^.

Tuberculosis (TB) caused by *Mycobacterium tuberculosis* remains a formidable threat to global public health challenge ^[2]^, with approximately 10 million new cases and and 1.5 million deaths globally reported in 2024^[3,4]^.The emergence of drug-resistant and multi-drug resistant (MDR) TB strains exacerbates this crisis, ^[5–8]^, underscoring the urgent need for novel diagnostics ^[9,10]^ and treatments are imperative to tackle the MDR-TB ^[11]^, especially new drug targets ^[12]^. *M. tuberculosis* combats host-derived oxidative stress ^[13–15]^ certain therapeutics^[16,17]^ through enzymes like catalase, superoxide dismutase, and peroxidase.

DNA methylation, an epigenetic modification mediated by DNA methyltransferases, regulates essential cellular processes including transcription, chromosome stability, DNA repair, transposition, drug resistance, and host-pathogen interactions ^[18–21]^. Differential methylation patterns are associated with mycobacterial virulence^[22,23]^. During Mtb infection, host genomic DNA methylation status is alerted ^[24,25]^, and influences immune responses ^[26]^. Mutations or dysregulation of bacterial DNA methyltransferase is impact survival^[27]^ and virulence ^[18,20,28,29]^. In Mtb, transcription factor binding can shield DNA regions from methylation ^[30]^ and Dam methylation typically represses instead of activating transcription ^[31]^. Mtb*-*infected macrophages exhibit differential methylation, particularly in genes involved in immune responses and chromatin remodeling ^[32]^. The methylation levels of key vitamin D pathway genes ^[33]^ and *Alu* elements^[34]^ correlated with the TB susceptibility, highlighting the significance of DNA methylation in Mtb-host interplay ^[35,36]^.

The Mtb genome encodes at least four DNA methyltransferases: two predicted (Rv3263, Rv3204; Mycobrowser) and two experimentally confirmed (Rv2966c, Rv2258c). Rv3263 is an N⁶-methyladenine (m⁶A) methyltransferase regulating gene expression and survival under hypoxia ^[27]^. Rv2258c, an S-adenosyl-L-methionine (SAM)-dependent methyltransferase, may contribute to drug resistance ^[37]^. The function of Rv3204 remains largely unexplored, beyond the observation that SAM is not its methyl donor^[32]^. While ROS and DNA methylation have been studied independently in mycobacteria, their interplay is unknown. Links between ROS, methylation, and processes like aging, ^[38,39]^ and cancer exist in eukaryotes. This study is the first to establish a connection between ROS and m^4^C DNA methylation in mycobacteria, demonstrating that Mtb Rv3204 is an m^4^C methyltransferase essential for managing oxidative stress and maintaining genomic integrity.

ROS or DNA methylation was separately studied in mycobacteria, but their interplay remains elusive. Previous studies have shown the relationship between the human aging process and ROS and methylation in cancer cells ^[40]^. In this study, we firstly established a link between ROS and m4C DNA methylation in *M. smegmatis*. We found that *M. tuberculosis* Rv3204 is an m4C DNA methyltransferase, Rv3204 homolog deleted *M.smegmatis* mutants accumulated higher amount of ROS, the survival rate of the deletion mutants was significantly decreased upon oxidative stress, the transcription of genes involved in DNA repair was also down-regulated. The result suggested that m4C DNA methyltransferase Rv3204 plays a role in *Mycobacterium* survival upon oxidative stress.

## Methods

### Construction of recombinant *M. smegmatis*

*Rv3204* was cloned by using the *M. tuberculosis* H37Rv genomic DNA and specific primers, PCR product was purified by using TIANquick Midi Purification Kit (Tiangen Biotechnology, Beijing, China), then ligated to plasmid pALACE directly. The recombinant plasmid Rv3204-pALACE and pALACE were electroporated into *M. smegmatis* mc^2^ 155, positive clones were verified by colony PCR.

### Construction of Ms_1939 deletion strain and complementation strain

An in-frame deletion of the gene *Ms_1939* (the homolog of *M. tuberculosis Rv3204*), two DNA fragments, comprising 800 bp, were amplified from the genome of *M. tuberculosis* H37Rv using specific primers, and cloned into the hygromycin excisable cassette. The resulting DNA fragment was purified and introduced into *M. smegmatis* derivative containing pJV53, a replicative plasmid expressing two phage recombinases and conferring kanamycin resistance. Three hygromycin-resistant colonies were isolated and tested by PCR for the correct integration of the excisable cassette into the chromosome. Subsequently, they were grown for 2 generations in the absence of hygromycin and kanamycin to allow the loss of hygromycin cassette and pJV53. One of them was analyzed in parallel with a colony of the wild-type parental strain by PCR with primers flanking the region used for the recombination. BGI sequencing confirmed the successful construction of the deletion mutant *M. smegmatis* (ΔMs_1939). Complementation was tested by amplifying the target gene *Rv3204* and cloning it into an integrating plasmid vector, pALACE.

### Strains and growth conditions

Bacterial strain *M. smegmatis* mc^2^ 155 were grown in 7H9 liquid medium supplemented with 0.05% (v/v) Tween 80 and 0.5% (w/v) glucose. Bacteria were cultured at 37℃ in shaker.

### Library construction for RNA-seq and sequencing procedures

Total RNA was isolated using RNeasy mini kit (Qiagen, Germany). The poly-A containing mRNA molecules were purified using poly-T oligo-attached magnetic beads. Following purification, the mRNA is fragmented into small pieces using divalent cations under 95℃ for 8 min. The cleaved RNA fragments are copied into first strand cDNA using reverse transcriptase and random primers. This is followed by second strand cDNA synthesis using DNA Polymerase I and RNase H. These cDNA fragments then go through an end repair process, the addition of a single ‘A’ base, and then ligation of the adapters.

The products are then purified and enriched with PCR to create the final cDNA library. Purified libraries were quantified by Qubit® 2.0 Fluorometer (Life Technologies, USA) and validated by Agilent 2100 bioanalyzer (Agilent Technologies, USA) to confirm the insert size and calculate the mole concentration. Cluster was generated by cBot with the library diluted to 10 pM and then were sequenced on the Illumina HiSeq 2500 (Illumina, USA). The library construction and sequencing was performed at Shanghai Biotechnology Corporation.

### Real-time PCR

Wild-type, ΔMs_1939 and ΔMs_1939 :: Rv3204 were cultured till OD_600_ over 1.0, bacterial was pelleted centrifugation at 8,000 rpm for 10 minutes. Total RNA was isolated by using TRIzol® reagent, DNA is digested by RNase-free DNase, then purified by RNAclean Tiangen kit (*Tiangen*, Beijing, China), cDNA is synthesized by using PrimeScript RT Reagent Kit (*Tiangen*, Beijing, China) with gDNA Eraser. RT-PCR Tiangen kit was used. Amplifications were carried out in reaction conditions as follows: 95℃ for 5min and a total of 40 cycles at 95℃ for 30 s, 60℃ for 30 s and 72℃ for 30 s. Gene expression levels were normalized to the levels of *sigA* gene transcription. All the gene specific primers used for RT-PCR were listed in the Supplementary Table 3.

### Growth curve

For growth curve measuring,Wild-type, ΔMs_1939 and ΔMs_1939 :: Rv3204 were cultured in 7H9 medium containing 0.1% (v/v) Tween 80, 1% (w/v) glucose and 1% (v/v) acetamide. OD_600_ was measured in every 3 hours for 2-3 days.

### H_2_O_2_ stress

Wild-type, ΔMs_1939 and ΔMs_1939 :: Rv3204 were cultured in 7H9 medium containing 0.1% (v/v) Tween 80, 1% (w/v) glucose and 1% (v/v) acetamide, 20mM H_2_O_2_ was added when OD_600_ =0.6, OD_600_ was measured in every 3 hours.

### Antibiotic stress

For growth curve measuring, strains were cultured in 7H9 medium containing 0.1% (v/v) Tween 80 and 1% (w/v) glucose, antibiotics (norfloxacin and ofloxacin) were added simultaneously at the concentration of 2μg/mL and 0.5μg/mL, respectively. OD_600_ was measured at an interval of every 3h.

### DNA methyltransferase inhibition

The inhibitor we used is 5-azacytidine (MedChemExpress Corp.). Wild-type, ΔMs_1939 and ΔMs_1939 :: Rv3204 was cultured till OD=0.6, then 20 mM H_2_O_2_ was added to 1mL liquid bacterial of one group, another group added nothing as a negative control, the final concentration of 5-azacytidine is 200 μM/mL, 500 μM/mL and 1 mM/mL, colonies was recorded at 0 h, 9 h, 24h, 48h, 72h, 96h.

### ROS measurement

Wild-type, ΔMs_1939 and ΔMs_1939::Rv3204 were cultured till OD_600_ =0.8-1.0, then pelleted centrifugation at 8,000 rpm for 10 min, precipitant was washed by 1×PBS for three times, then adjusted the OD_600_ to around 0.8, specific ROS-detecting fluorescent dye 2’,7’-dichlorodihydrofluorescein diacetate (DCFH-DA) was diluted with 1× PBS (1:100), 20μL dye was exposed to 180 μL bacterial at 37℃ for 30min, then transferred to 96-well microplates, 200μL for each well, the fluorescence intensity was measured by using fluorescence plate reader, excitation wavelength is 485nm and emission wavelength is 525nm.

### DNA damage analyses

A terminal deoxynucleotidyl transferase dUTP nick-end labeling (TUNEL) assay was performed to detect DNA damage by using the One-Step TUNEL Apoptosis Assay Kit (Beyotime, Jiangsu, China). Wild-type, ΔMs_1939 and ΔMs_1939 :: Rv3204 were cultured in 7H9 medium containing 0.1% (v/v) Tween 80, 1% (w/v) glucose and 1% (v/v) acetamide to OD_600_ = 0.8-1.0, with or without antibiotics (norfloxacin), bacteria were pelleted centrifugation at 8,000 rpm for 2 minutes and washed with pre-cooled 1×PBS for two times. The bacteria were resuspended with 500μL 1× PBS and then fixation solution was added (final concentration 2% PFA) at room temperature for 30 minutes. After 3 minutes of centrifugation at 9,000 rpm, precipitate was washed with 1×PBS, bacteria was centrifuged at 8,000 rpm for 3 minutes and then suspended with 1×PBS containing 0.3% Triton X-100, co-incubated at room temperature for 5 minutes. 50μL TUNEL reaction mixture was added after two times washing, then incubated for 60 minutes at 37℃ in the dark. Bacteria were washed for twice with 1×PBS and suspended with 400μL 1×PBS, the fluorescence intensity was measured by using flow cytometer, excitation wavelength is 450-500nm and emission wavelength is 515-565nm.

### Bioinformatic Analysis

The open reading frame and catalytic domain were confirmed using the DNAMAN bioinformatics software and NCBI-BLAST (https://blast.ncbi.nlm.nih.gov/Blast.cgi). And we analyzed the data for methylation and transcriptomics by R statistical software environment and R Programming language version 3.6.2.

### Statistical analysis

Data from at least three biological replicates were used to calculate means and standard deviation (SD) for graphing purposes. Statistical analysis employed the unpaired student’s t test, asterisks indicate statistically significant difference (*P < 0.05; **P < 0.01; ***P < 0.001).

## Results

### Rv3204 is a conserved DNA methyltransferase

(A) *M. tuberculosis* genome lacks Dam or Dcm homologs^[41]^. However, the presence of 6-methyladenine and 5-methylcytosine modifications in *M. tuberculosis* and *M. smegmatis* genomes suggests the alternative methyltransferases exist. *M. tuberculosis* genome has two predicted DNA methyltransferases, namely *Rv3263* and *Rv3204* (https://mycobrowser.epfl.ch/). Rv3263 (MamA) was a methyltransferase for N^6^-MdA^[27]^, the characteristics of Rv3204 remain poorly addressed. *Rv3204* is highly conserved across mycobacterial species, from non-pathogen to pathogen (Fig 1A). It encodes a 101 amino acid protein featuring an Ada-like DNA-binding domain (Fig 1B; NCBI Gene: 888132). Meanwhile, its *M. smegmatis* homolog, Ms_ 1939 shares the same active site and domain architecture (Fig 1C; NCBI Gene: 4536670), but lacks the SAM-binding motif (^51^YAGSG^55^), consistent with previous findings that SAM is not its methyl donor ^[42]^.

**Figure 1.**
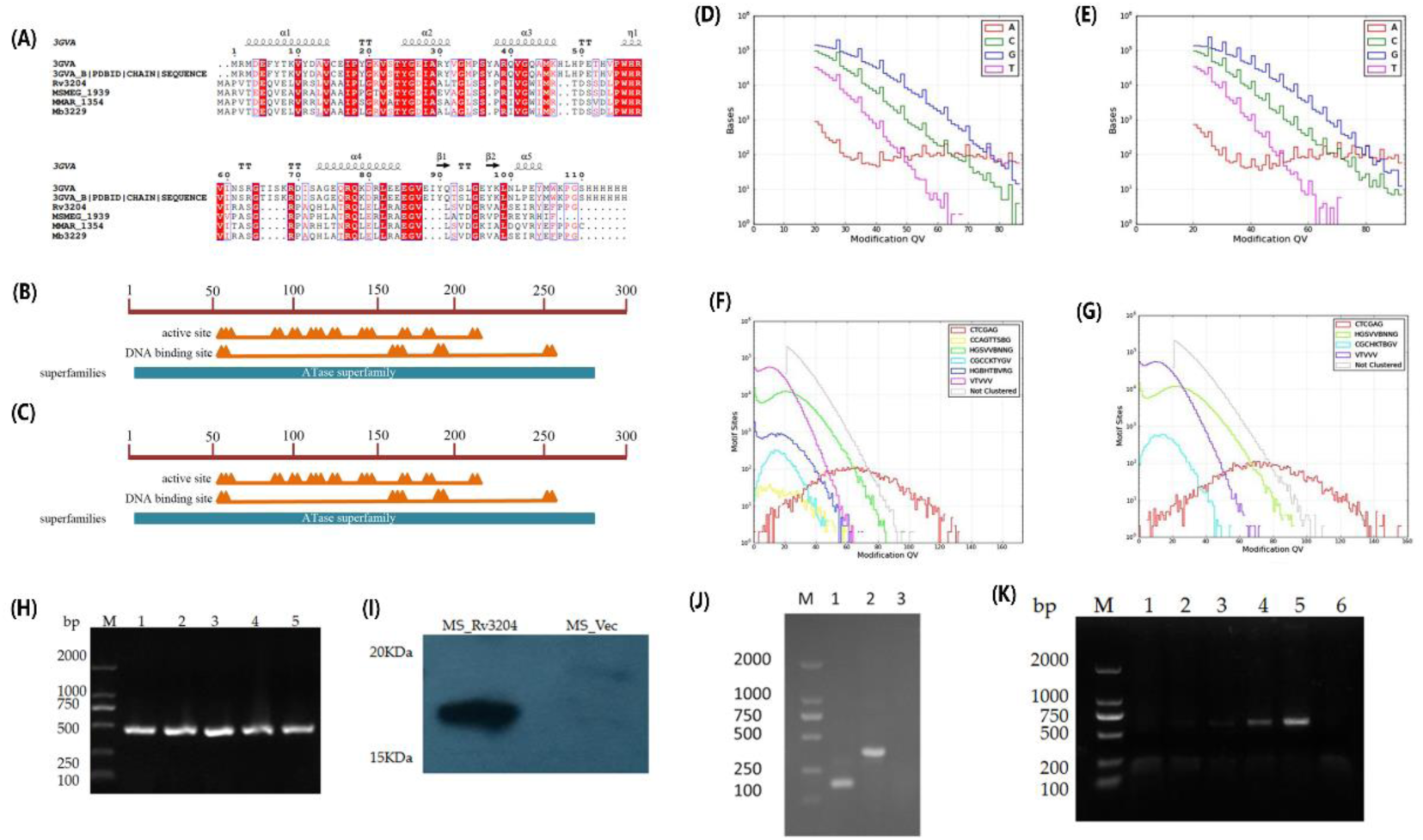
Conservation analysis of Rv3204, methylation alterations in ΔMs_1939, and heterologous expression validation (A) Sequence alignment of Rv3204 homologs: *M. tuberculosis* H37Rv Rv3204c; *M. bovis* Mb3229; *M. marinum* MMAR_1354; *M. smegmatis* Ms_1939 (3GVA: S. pombe ATL crystal structure reference). (B–C) Conserved domain analysis: (B) Rv3204; (C) Ms_1939. (D–E) Altered DNA methylation in ΔMs_1939: (D) Methylation levels of adenine, thymine, cytosine, and guanine in wild-type; (E) Significant cytosine methylation changes in ΔMs_1939. (F–G) Methylation motif comparison: (F) Motifs CCAGTTSBG and HGBHTBVRG in wild-type; (G) Loss of both motifs in ΔMs_1939. (H–K) Heterologous expression in *M. smegmatis*: (H) PCR amplification of Rv3204c (∼300 bp); (I) Western blot detection of His-tagged Rv3204 in recombinant strains; (J) ΔMs_1939 mutant construction (Lane 1: Wild-type; 2: ΔMs_1939; 3: Negative control); (K) Complementation strain ΔMs_1939::Rv3204 (Lane 5: Complemented strain; 6: Negative control; M: DL 2,000 DNA marker).

### Rv3204 is a possible m4C DNA methyltransferase

We successfully constructed a strain over-expressing Rv3204c protein in *M.smegmatis* which has high homology with *M.tuberculosis*, and successfully knocked out Rv3204c homologous gene Ms_1939 in *M.smegmatis* (Ms_1939, ΔMs_1939). At the same time, we also constructed the complementary strain which the Rv3204c is overexpressed in the knockout strain (ΔMs_1939::Rv3204) ‘Fig 1 H-K’. To define Rv3204’s methyltransferase specificity, we performed single-molecule real-time (SMRT®) sequencing to map methylomes at single-base resolution in the wild-type (WT) and ΔMs_1939 strains. Compared to WT, the ΔMs_1939 genome exhibited a substantial loss of cytosine methylation (Fig. 1 D, E). Motif analysis revealed the absence of two methylation motifs in the mutant: CCAGTTSBG (modification type: m^4^C) and HGBHTBVRG (modification type uncertain) (Fig. 1F, G). These data confirm Rv3204 as an m^4^C DNA methyltransferase specific for the CCAGTTSBG motif. Notably, another m^4^C motif (CGCCKTYGV) was detected in both strains, but its frequency was significantly higher in ΔMs_1939 (955 vs 564 sites in WT), the cause of which warrants further investigation.

### Altered methylation and expression of ROS scavenging and DNA repair genes in ΔMs_1939

Integrated DNA-seq and RNA-seq analyses revealed significant differences in the expression of ROS detoxification and DNA repair genes between WT and ΔMs_1939. Notably, genes belonging to the Mut DNA repair family exhibited differential expression (Fig. 2F) and methylation (Fig. 2G), suggesting potential epigenetic regulation. Quantitative RT-PCR (qRT-PCR) confirmed that under basal conditions, transcription levels of key ROS scavenging genes (*katG, sodA1, sodA2, sodC*) were comparable across strains (Fig. 3B). However, upon oxidative stress (implied by context), the mutant failed to upregulate these genes effectively (linking back to Abstract/Fig 3D phenotype).

**Figure 2.**
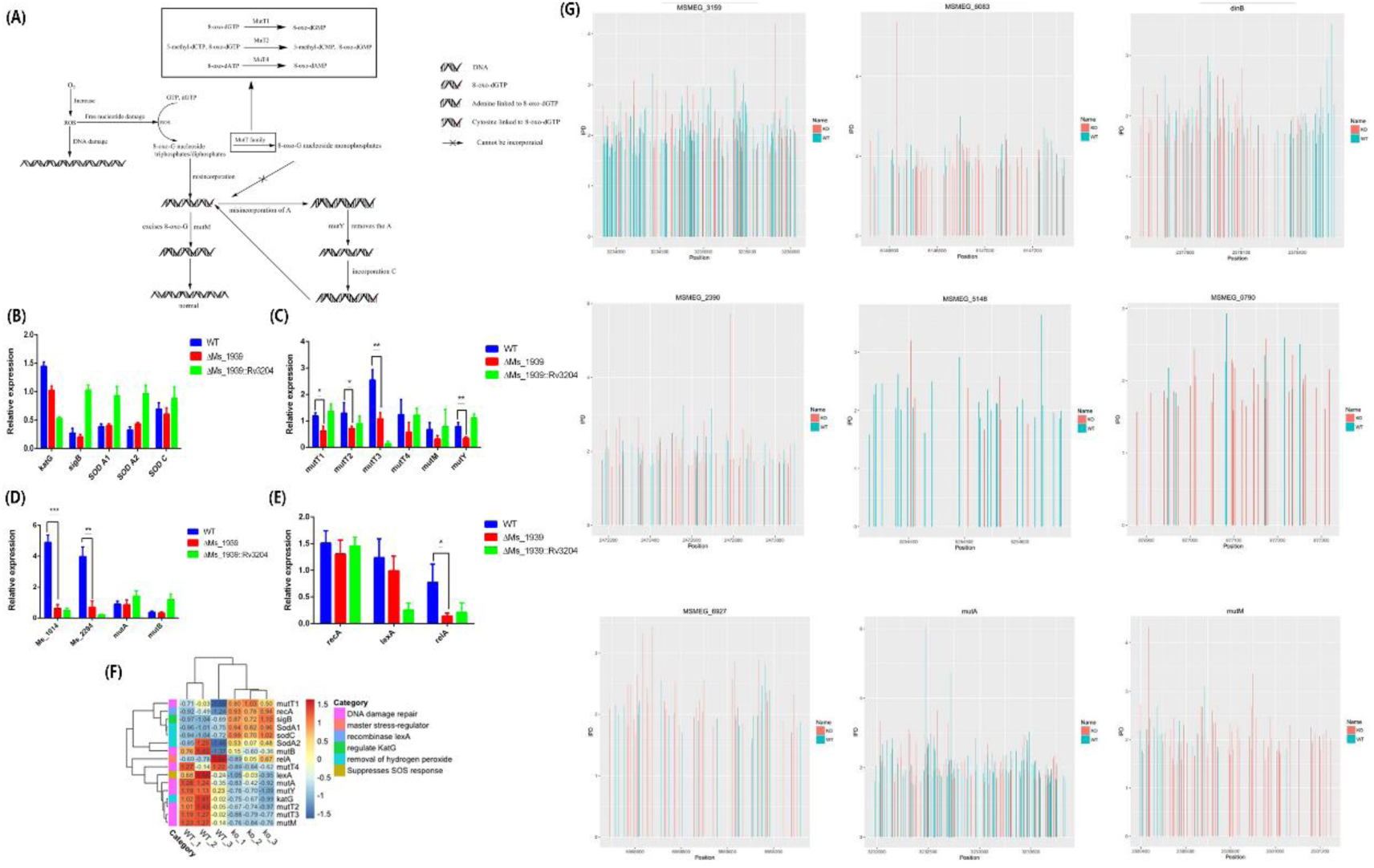
Differential gene expression, qRT-PCR validation, and methylation analysis in WT versus ΔMs_1939. (A) Heatmap of differentially transcribed genes between WT and ΔMs_1939. Up-regulated genes (red) and down-regulated genes (blue) are shown. Data represent three biological replicates. (B) Schematic of Mut family proteins in ROS-induced DNA damage repair. Key: ROS, reactive oxygen species; GTP, guanosine triphosphate; dGTP, 2’-deoxyguanosine 5’-triphosphate; mut family, DNA mutation repair genes; A, adenine; C, cytosine; 8-oxo-G, 8-oxo-7,8-dihydroguanine. (C) qRT-PCR analysis of ROS scavenging gene expression. (D-F) qRT-PCR analysis of mut family and DNA damage-related genes: (D) mutT1, (E) mutT2, (F) mutT3 (function unknown). (G) Differential methylation of Mut family motifs. Bar height indicates IPD (inter-pulse duration) ratio. Blue: WT methylation; red: ΔMs_1939 methylation. Data in (C-F) shown as mean ±SD of triplicate wells. Statistical analysis by two-sided Student’s t-test: *P < 0.05, **P < 0.01, ***P < 0.001. All qRT-PCR analyses performed using GraphPad Prism 6.0.

**Figure 3.**
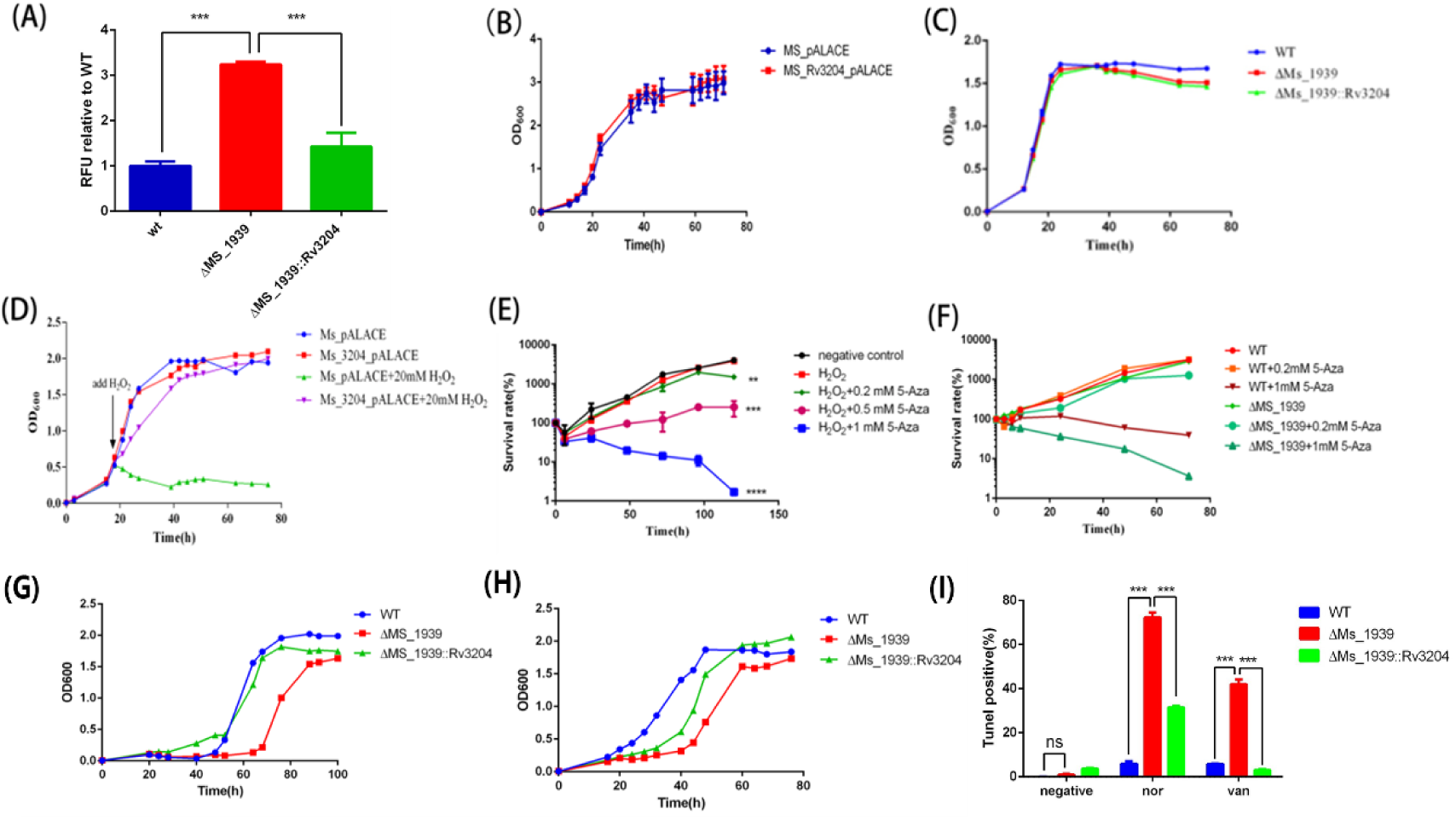
Phenotypic characterization of ΔMs_1939: ROS accumulation, stress tolerance, and DNA damage. (A) Intracellular ROS levels in ΔMs_1939 versus wild-type (WT). (B) Growth comparison: Control (Ms_pALACE) vs. Rv3204-expressing recombinant (Ms_Rv3204_pALACE). (C) Growth curves under normal conditions: WT, ΔMs_1939, and complemented strain (ΔMs_1939::Rv3204c). (D) H₂O₂ tolerance (20 mM): Survival of control vector (Ms_pALACE) vs. Rv3204-expressing strain (Ms_pALACE_Rv3204). (E) 5-azacytidine inhibition: Concentration-dependent abolition of H₂O₂ tolerance in Ms_Rv3204_pALACE. (F) Differential 5-azacytidine sensitivity: WT vs. ΔMs_1939 at indicated concentrations. (G-H) Fluoroquinolone stress in 7H9 medium at 37°C: (G) Growth under norfloxacin; (H) Growth under ofloxacin. (Both: WT, ΔMs_1939, complemented strain) (I) DNA damage levels with/without antibiotics: norfloxacin (2 μg/mL), vancomycin (20 μg/mL). Strains grown to OD₆₀₀ = 0.8-1.0. All data represent mean ±SD of triplicate wells. Statistical significance determined by two-sided Student’s t-test: *P < 0.05, **P < 0.01, ***P < 0.001. All experiments reproduced three times with consistent results.

Phagocytes eliminate pathogens via ROS production (41), countered by bacterial DNA repair systems (42). qRT-PCR analysis of DNA damage response genes (Supplementary Table 2) revealed significant downregulation in ΔMs_1939 compared to WT (Fig. 3C, D, E). Affected genes included *mutT1, mutT2, mutT3, mutY, Ms_1014, Ms_2294*, and the stress regulator *relA*. This downregulation suggests impaired DNA repair capacity in the mutant. However, potential post-transcriptional regulation affecting protein stability and activity should be considered.

Collectively, these data indicate that Rv3204 deficiency compromises the transcriptional response necessary for ROS detoxification and DNA repair, aligning with the observed hypersensitivity of ΔMs_1939 to oxidative stress.

### The intracellular accumulation of ROS was significantly increased in deletion mutant ΔMs_1939

To determine if altered H₂O₂ susceptibility stemmed from differences in ROS handling, we measured intracellular ROS levels using DCFH-DA fluorescence. The ΔMs_1939 mutant exhibited significantly higher basal levels of intracellular ROS compared to WT and the complemented strain (ΔMs_1939::Rv3204) (Fig. 3A), indicating that Rv3204 is essential for maintaining ROS homeostasis.

### Rv3204 confers tolerance to hydrogen peroxide

To assess the role of Rv3204 in oxidative stress survival, we monitored bacterial growth. Under standard aerobic conditions, strains overexpressing Rv3204 (Ms_Rv3204_pALACE) grew similarly to the vector control (Ms_pALACE) (Fig. 3B), and no growth difference was observed among WT, ΔMs_1939, and ΔMs_1939::Rv3204 (Fig. 3C). However, upon exposure to 20 mM H₂O₂, the Rv3204-overexpressing strain (Ms_Rv3204_pALACE) exhibited robust growth, while the vector control (Ms_pALACE) failed to grow (Fig. 3D), demonstrating that Rv3204 enhances H₂O₂ tolerance.

To confirm this phenotype depends on Rv3204’s methyltransferase activity, we treated the Rv3204-overexpressing strain with the DNA methyltransferase inhibitor 5-azacytidine. Increasing concentrations of 5-azacytidine progressively abolished the H₂O₂ tolerance conferred by Rv3204 overexpression (Fig. 3E). Furthermore, under inhibitor treatment, WT showed greater inherent tolerance than the ΔMs_1939 mutant (Fig. 3F), consistent with Rv3204’s protective role being dependent on its enzymatic activity.

### ΔMs_1939 exhibits impaired growth under fluoroquinolone stress

ROS contribute significantly to the bactericidal activity of fluoroquinolones, key second-line TB drugs. Reduced ROS accumulation can paradoxically enhance bacterial survival at high drug concentrations. We assessed the growth of WT, ΔMs_1939, and ΔMs_1939::Rv3204 under sub-inhibitory concentrations of norfloxacin or ofloxacin. The ΔMs_1939 mutant, which accumulates more ROS (Fig. 3A), displayed significantly slower growth compared to WT and the complemented strain upon exposure to both antibiotics (Fig. 3G, H), indicating heightened susceptibility.

### Increased DNA damage susceptibility in ΔMs_1939

Given the downregulation of DNA repair genes in ΔMs_1939 (Fig. 2C-E), we hypothesized increased DNA damage susceptibility. Using TUNEL (TdT-mediated dUTP Nick-End Labeling) assays, we quantified DNA damage in WT, ΔMs_1939, and ΔMs_1939::Rv3204 strains, with or without antibiotic exposure. The ΔMs_1939 mutant exhibited significantly higher levels of DNA damage, both basally and upon treatment with norfloxacin (2 μg/mL) or vancomycin (20 μg/mL), compared to WT and the complemented strain (Fig. 3I). This confirms that loss of Rv3204 compromises genomic integrity and renders the bacterium more vulnerable to DNA-damaging agents.

## Discussion

The role of bacterial epigenetics, particularly DNA methylation, is an area of intense investigation. This study identifies *M. tuberculosis* Rv3204 as an m^4^C DNA methyltransferase. Deletion of its homolog in *M. smegmatis* (ΔMs_1939) increased susceptibility to oxidative stress, elevated intracellular ROS levels, impaired transcriptional upregulation of ROS detoxification and DNA repair genes, heightened DNA damage, and increased sensitivity to fluoroquinolones. These findings establish a novel link between m^4^C DNA methylation and ROS management in mycobacteria, revealing a role for epigenetic regulation in bacterial survival under oxidative stress. The observed phenotypic differences likely stem from combined epigenetic and transcriptomic alterations, positioning Rv3204 as a promising target for novel anti-TB strategies aimed at disrupting ROS homeostasis.

We characterized the first m4C DNA methyltransferase of *M.tuberculosis* by using the SMRT sequencing which permits the genome-wide recognition of methylated bases in bacteria. The motif of the m4C DNA methyltransferase Rv3204 is CCAGTTSBG. The amino acid identity between Rv3204 and Ms_1939 is up to 77.23%, suggesting that the methylation motif might also be conserved in the *M.tuberculosis*. This is the first to reveal all three types of methylation in *Mycobacterium* using SMRT sequencing. 5-methylcytosine (m5C) methylation was reported in *M.tuberculosis*^[42]^. m4C was previously documented in *Helicobacter pylori*^[43]^, but the motif is not CCAGTTSBG. The motif discrepancy between *H.pylori* and *Mycobacterium* might be due to their GC content difference. DNA methylomes revealed that two motifs (CCAGTTSBG and HGBHTBVRG) are absent in the DNA methyltransferase deletion mutant ΔMs_1939. The modification type of CCAGTTSBG is m4C, while the modification of HGBHTBVRG is uncertain, indicating that Rv3204 is an m4C methyltransferase. Another m4C methylation motif CGCCKTYGV was shared by both ΔMs_1939 and wild type, but the motif detected in deletion mutant is 955, twofold of the wild type which is 564. The underlying cause remains to be determined.

The ΔMs_1939 phenotype—increased ROS accumulation, failure to induce ROS detoxification genes, downregulation of DNA repair genes, heightened DNA damage, and increased antibiotic susceptibility—can be interpreted through several mechanisms. The methyltransferase may directly regulate the transcription of key defense genes via promoter methylation, indirectly influence regulators of the oxidative stress response, or impact global genomic stability. The observed accumulation of ROS and impaired transcriptional response in the mutant provide a plausible explanation for its increased vulnerability to DNA damage, consistent with established mechanisms ^[44]^.

The catalase activity was correlated with pathogen virulence ^[45]^. Nucleotide oxidation under ROS stress does not affect DNA replication but resulting in mutations because of base mismatch^[46]^, bacterial survival will be compromised. The oxidative damage can be DNA deoxyribose resulting in DNA strand breaks or oxidize the free guanine into oxidized guanine, which is ambiguous when pairing with cytosine and adenine^[47]^. Among the four bases that make up DNA, guanine is more vulnerable to oxidation than other three bases due to its lower reduction potential^[48]^. Mis-incorporation of 8-oxo-dGTP in DNA can be detrimental to genomic integrity. Adenine methylation and cytosine methylation is also related to overall background genomic mutation rate^[49,50]^.

Mut proteins play an important role in bacterial survival under oxidative stress^[46]^, mutT proteins are sanitization enzymes which can hydrolyze nucleotides such as 8-oxo-dGTP to their respective monophosphate products^[51]^, mutM recognizes and excises oxidized purines^[52]^, mutY excises mis-incorporated A against 8-oxoG^[53]^. The decrease of *mut*T expression in ΔMs_1939 indicates a decrease in the ability of the bacteria to resist oxidative damage, that is, the loss of methyltransferase may directly or indirectly affect the ability of the bacteria to repair DNA. We discovered that the gene related to DNA repair was differently methylated in WT and ΔMs_1939, which may be associated with the low expression of *mut* family genes in ΔMs_1939.

The conserved domain of Rv3204 belongs to the ATase superfamily, and the product of Ms_1939, the homologous gene in *M. smegmatis*, is 6-O-methylguanine-DNA methyltransferase. Previous studies have shown that 6-O-methylguanine-DNA methyltransferase (MGMT, also known as ATase, AGT, AGAT) can remove temozolomide-induced DNA alkylation to prevent or even reverse DNA damage as a DNA repair protein^[54,55]^. Rv3204 might also have DNA repair capability, reconciled with some ΔMs_1939 phenotypes observed.

In summary, we have established Rv3204 as a novel m^4^C DNA methyltransferase essential for maintaining ROS homeostasis and genomic integrity in mycobacteria. Its critical role in survival under oxidative and antibiotic stress positions Rv3204 as a compelling candidate for therapeutic intervention in tuberculosis.

## Acknowledgements

We thank Shanghai Biotechnology Corporation for metabolome and transcriptome technical support.

## Supplementary material

**Supplement Table 1.** Genes involved in ROS clearance.

| Gene number | Homolog | Product | function |
| --- | --- | --- | --- |
| **Rv2710** | Ms_2752 | RNA polymerase sigma factor SigB | regulate KatG (Rv1908c) and the heat-shock response. |
| **Rv1098c** | Ms_6384 | Catalase-peroxidase-peroxynitritase T KatG | Participate in the removal of hydrogen peroxide |
| **Rv3846** | Ms_6427 | Superoxide dismutase [Fe] SodA1 | Destroys radicals normally produced within the cells and toxic |
| **Rv3846** | Ms_6636 | Superoxide dismutase [Fe] SodA2 | Destroys radicals normally produced within the cells and toxic |
| **Rv0432** | Ms_0835 | Periplasmic superoxide dismutase [Cu-Zn] SodC | Destroys radicals normally produced within the cells and toxic |

**Supplementary Table 2.** Genes related to DNA repair Gene number Homolog Name Product.

| Gene number | Homolog | Name | Product |
| --- | --- | --- | --- |
| Rv1160 | Ms_5148 | *mutT2* | mutator protein MutT2 (7,8-dihydro-8-oxoguanine-triphosphatase) (8-oxo-dGTPase) |
| Rv1492 | Ms_3158 | *mutA* | methylnalonyl-CoA mutase small subunit MutA (MCM) |
| Rv1493 | Ms_3159 | *mutB* | methylnalonyl-CoA mutase large subunit MutB (MCM) |
| Rv0413 | Ms_0790 | *mutT3* | mutator protein MutT3 (7,8-dihydro-8-oxoguanine-triphosphatase) (8-oxo-dGTPase) (dGTP pyrophosphohydrolase) |
| Rv2985 | Ms_2390 | *mutT1* | hydrolase MutT1 |
| Rv3589 | Ms_6083 | *mutY* | adenine glycosylase MutY |
| Rv3908 | Ms_6927 | *mutT4* | mutator protein MutT4 |
| Rv2924c | Ms_2419 | *mutM* | formamidopyrimidine-DNA glycosylase Fpg<br>(FAPY-DNA glycosylase) |
| Ms_1014 |  | *dinB1* | DNA polymerase IV |
| Ms_2294 |  | *dinB2* | DNA polymerase IV |

**Supplementary Table 3.**
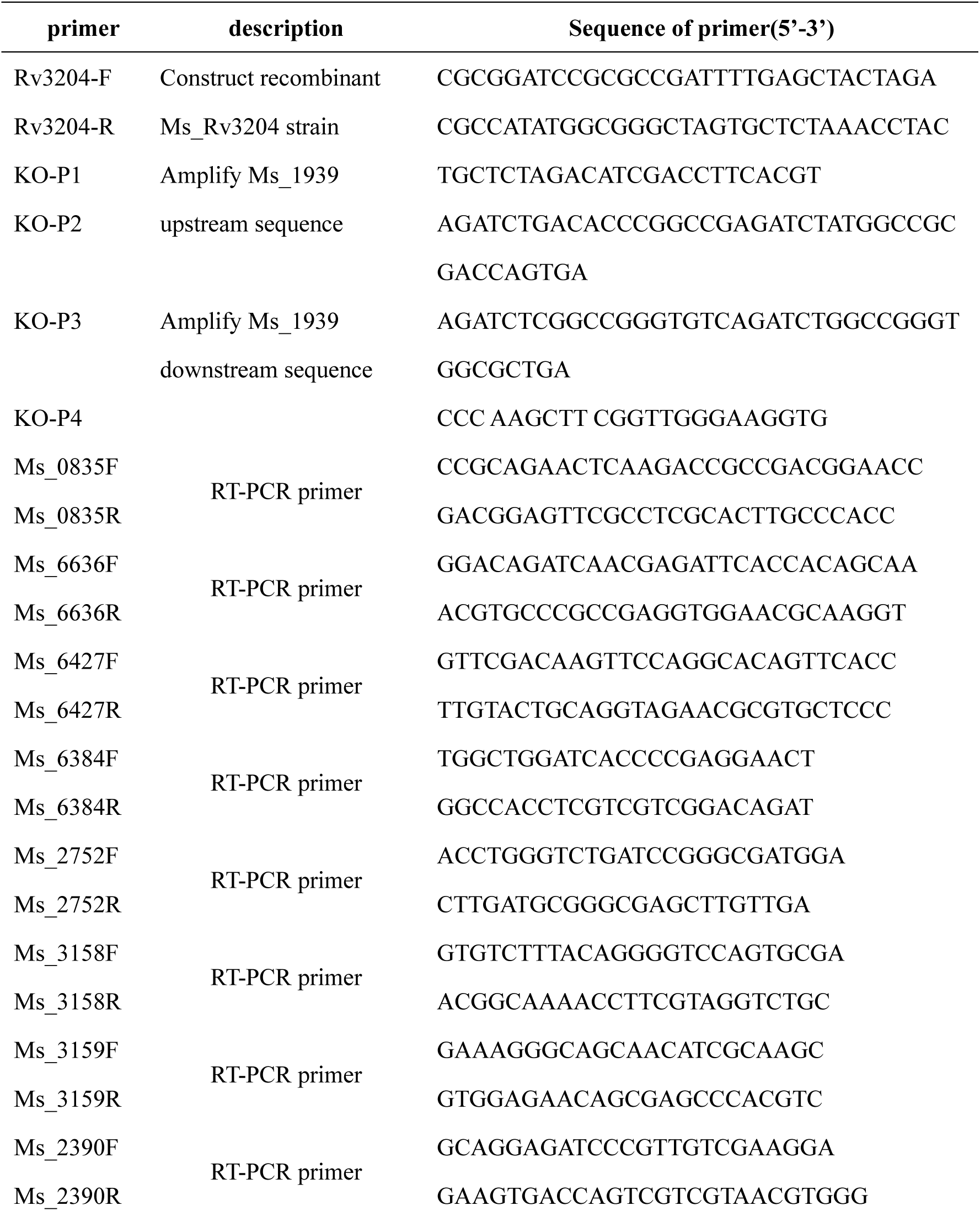
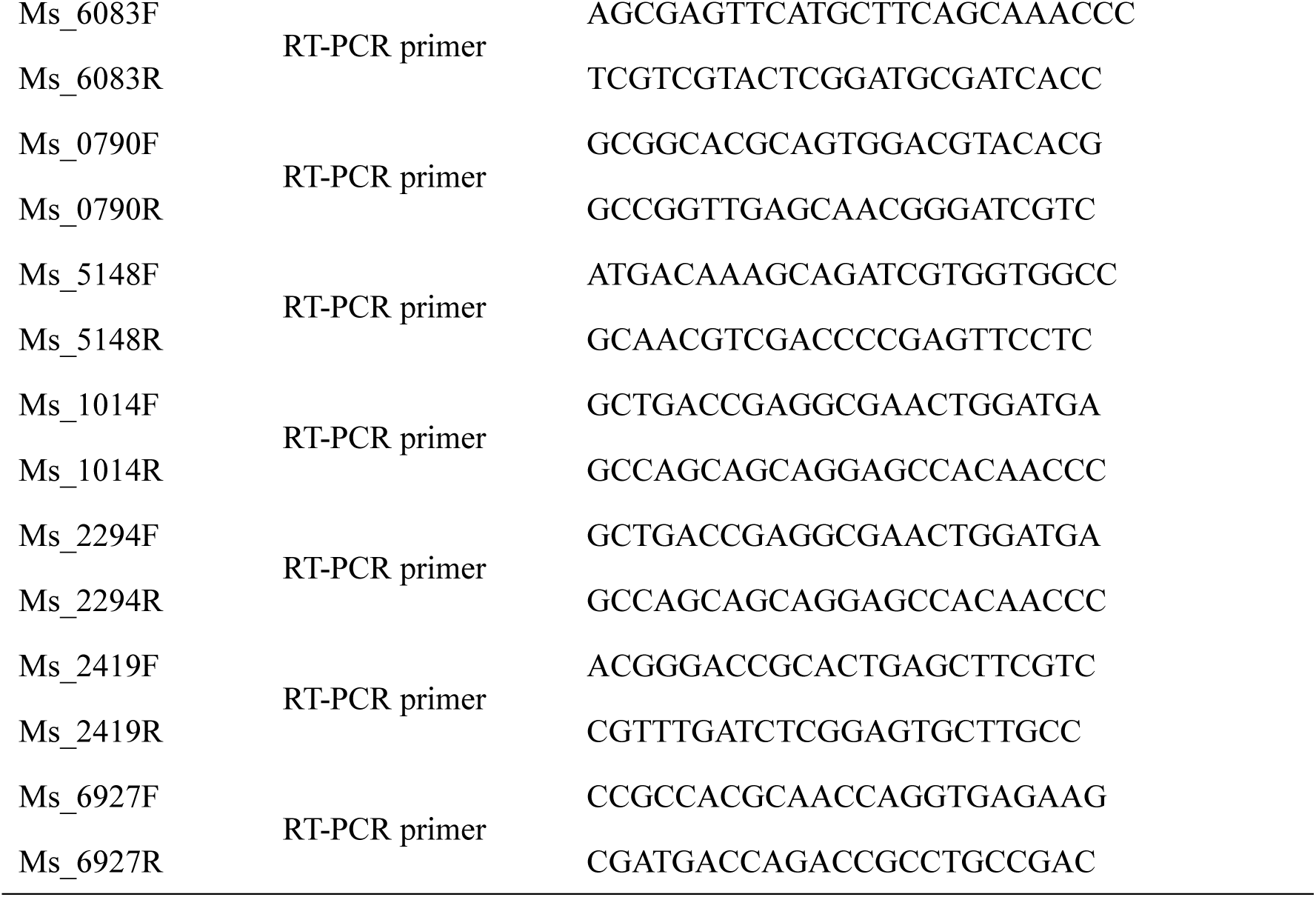
Primers used in this study primer description Sequence of primer(5’-3’)

